# Quantitative site-specific glycoproteomics by ZenoTOF reveals glyco-signatures for breast cancer diagnosis

**DOI:** 10.1101/2024.09.08.611557

**Authors:** Yi Yang, Dan Zhao, Ji Luo, Ling Lin, Yuxiang Lin, Baozhen Shan, Hongxu Chen, Liang Qiao

**Author notes:** **Corresponding Author** Correspondence should be addressed to Dr. Baozhen Shan, Dr. Hongxu Chen or Prof. Dr. Liang Qiao. These authors contributed equally: Yi Yang, Dan Zhao, and Ji Luo.

## Abstract

Intact glycopeptide characterization by mass spectrometry has proven a versatile tool for site-specific glycoproteomics analysis and biomarker screening. Here, we present a method using the ZenoTOF instrument with optimized fragmentation for intact glycopeptide identification and demonstrate its ability to analyze large-cohort glycoproteomes. From 124 clinical serum samples of breast cancer, non-cancerous diseases, and non-disease controls, a total of 6901 unique site-specific glycans on 807 glycosites of proteins were detected. Much more differences of glycoproteome were observed in breast diseases than the proteome. By employing machine learning, 15 site-specific glycans were determined as potential glyco-signatures in detecting breast cancer. The results demonstrate that our method provides a powerful tool in glycoproteomic analyses for biomarker discovery studies.

---

Protein glycosylation is a critical post-translational modification (PTM) that mediate diverse biological functions^1-4^. Aberrant glycosylation is well associated with diseases, such as inflammation and tumors^5-7^. Thus, glycosylation is an essential target for disease-related biomarker discovery. Glycosylation is chemically and biosynthetically diverse not only in the occupancy of individual sites of glycosylation (macroheterogeneity), but also in the composition and structure of the attached glycans at specific sites (microheterogeneity). In respect of these heterogeneities, intact glycopeptide characterization enables the analysis of site-specific glycosylation on a proteome-wide scale^8^, which is an imperative but challenging component to modern glycoproteomic studies.

Liquid chromatography–tandem mass spectrometry (LC-MS/MS) is the method of choice widely used in glycoproteomics, which characterizes glycopeptides by the fragmentation of precursor glycopeptide ions to analyze the resultant product ions^9, 10^. Various dissociation methods, such as higher-energy collisional dissociation (HCD) with different collisional energy (CE) settings, have been developed and optimized to characterize glycopeptides^11^. For N-glycopeptides, HCD with stepped CE on Orbitrap hybrid or tribrid mass spectrometers can generate comprehensive fragments for an intact glycopeptide, providing information of both the underlying peptide backbone and the attached glycan in a single MS/MS spectrum^11^. This method has been widely used for N-glycoproteomic studies, including drug resistance of tumors^12^, biomarker discovery for disease diagnosis^13, 14^, and host–virus interactions^15, 16^.

While most of the glycoproteomics studies used Orbitrap-based instruments, quadrupole time-of-flight (Q-TOF) systems provide an alternative with high acquisition rates and ion usage. Beam-type collision induced dissociation (CID) on Q-TOF systems can produce similar MS/MS spectra of peptides to Orbitrap-specific HCD^17^. With recent advances in instrumental technology, a new model of Q-TOF system named ZenoTOF employs a Zeno trap to increase the duty cycle and boost the MS/MS sensitivity, facilitating higher identification and quantification performance in proteomics analyses^18^. ZenoTOF is a promising choice for glycoproteomics, while it calls for the development of glycopeptide characterization methods for accurate and in-depth analysis of complex glyco-proteome samples.

Herein, we present a method using ZenoTOF for analyzing glycoproteomes from a large cohort of serum samples to screen disease-related signatures. The CID parameters were tuned towards a dynamic and stepped CE that was optimal for intact glycopeptide fragmentation and identification. The method was applied to analyze the glycoproteomes of 124 clinical serum samples of breast cancer, non-cancerous diseases, and non-disease controls. A total of 6901 unique site-specific glycans from 807 glycosites of proteins were detected from the cohorts. Proteomic analysis of the samples was conducted in parallel and differential abundance analyses demonstrated that the glycoproteome was more variable in breast diseases than the proteome. By employing machine learning, 15 site-specific glycans were determined as potential glyco-signatures in detecting breast cancer. The results demonstrated that our method provides a powerful tool in glycoproteomic analyses for biomarker discovery studies.

## EXPERIMENTAL SECTION

### Clinical sample collection

Clinical serum samples were collected from 40 patients diagnosed with breast malignant tumor (MT), 43 patients diagnosed with benign breast lesions (BL), and 41 control volunteers without any history of breast diseases (**Table S1**). Samples were collected prior to any therapeutic interventions, spanning the years 2017 to 2022 in the Fujian Medical University Union Hospital. Explicit written informed consent was acquired from all subjects, affirming the exclusive usage of these biological materials for research purposes. The study protocol received approval from the Ethics Committee of Fujian Medical University Union Hospital (Fuzhou, China) (Approval No. 20200321) and adhered strictly to all pertinent laws and regulations of China governing biomedical research.

### Sample preparation and MS analysis

Proteins from individual human sera were denatured by urine, reduced by tris(2-carboxyethyl)phosphine, alkylated by 2-chloroacetamide, digested by trypsin, and desalted by C18 cartridges. Glycopeptides were enriched using hydrophilic interaction liquid chromatography.

The proteome and glycoproteome samples were submitted for microflow LC-MS/MS analysis using a ZenoTOF 7600 mass spectrometer. For glycoproteomic analysis, data-dependent acquisition (DDA) was used for MS/MS analysis. For benchmarking purpose, a pooled glycopeptide sample was analyzed using different CID parameters. The clinical glyco-proteome samples were analyzed using CID with dynamic CE. For the clinical proteome samples, data-independent acquisition (DIA) was used.

Details are provided in the Supporting Information **Method S1**.

### Data analysis and bioinformatics

For the benchmarking glycoproteome samples, GlycanFinder^19^, PEAKS Studio^19^, and FragPipe^20, 21^ were used for glycopeptide identification. For the clinical glycoproteome samples, GlycanFinder was used for glycopeptide identification and quantification. For the clinical proteome samples, Spectronaut^22^ was used for protein identification and quantification without the use of a spectral library (i.e., the directDIA+ workflow).

Bioinformatic analysis in this study included differential expression analysis by limma^23^ and volcano3D^24^, enrichment analysis by clusterProfiler^25^, machine learning-based biomarker screening by MetaboAnalystR^26^. Multiple packages, including VennDiagram^27^, ComplexHeatmap^28^, and volcano3D^24^, were used for data visualization.

Details are provided in the Supporting Information **Method S2**.

## RESULTS

### Characterizing intact *N*-glycopeptides by ZenoTOF

We established a workflow of MS acquisition and data analysis for intact glycopeptide identification using the Zeno-TOF instrument. We surveyed the CID parameters to optimize conditions for glycopeptide fragmentation and identification. The dynamic collisional energy (CE) was chosen where the CE center points automatically adjusted according to the mass-to-charge ratio (*m*/*z*) of the precursor ions, with a collisional energy spread (CES). The setting of CES was analogous to stepped CE for Orbitrap hybrid or tribrid mass spectrometers^11^, where sub-spectra were acquired at each CE (ramped from CE−CES to CE+CES) and summed into a single MS/MS spectrum during MS/MS acquisition.

We explored the MS informatics solutions for glycopeptide identification. We performed 3 technical replicates (repeated LC–MS/MS injections) to analyze the pooled serum sample using the dynamic CE. We benchmarked GlycanFinder 2.0^19^, PEAKS (Glycan module)^19^ and FragPipe (MSFragger-glyco)^20^ for glycopeptide identification. Other software tools that are commonly used for Orbitrap data, such as pGlyco^29^, Glyco-Decipher^30^, and StrucGP^31^, were not involved in this study due to their incompatibility with the ZenoTOF data. The results were reported with a glycopeptide false discovery rate (FDR) or q-value threshold of 1% using the quality control scheme implemented in each software. GlycanFinder 2.0 identified 1583±37 (mean ± standard deviation, sic passim) glycopeptides, 1539±28 site-specific glycans, and 232±3 glycosites per injection (**Figure S1a**), which was 30% more than PEAKS (1216±11 glycopeptides, 1196±15 site-specific glycans, and 176±2 glycosites) and 11%–19% more than FragPipe (1426±98 glycopeptides, 1304±82 site-specific glycans, and 195±6 glycosites). Accumulating the 3 replicate injections, 2494 glycopeptides, 2392 site-specific glycans, and 288 glycosites were identified totally by GlycanFinder 2.0, among which 35% glycopeptides, 36% site-specific glycans, and 62% glycosites were shared in all the replicates (**Figure S1b**). PEAKS and FragPipe resulted in 38% (702/1864 and 840/2189) glycopeptides, 39%–40% (721/1803 and 781/1990) site-specific glycans, and 68%–74% (143/210 and 166/224) glycosites shared in all the replicates. The data completeness level was consistent with our previous serum glycoproteomics study using an Orbitrap-based mass spectrometer^32^. Considering proteins and peptides shared in at least 2/3 replicates, GlycanFinder 2.0 resulted in a gain of 28% (1384 compared to 1083) glycopeptides and 27% (1351 compared to 1063) site-specific glycans, and 30% (229 compared to 176) glycosites than PEAKS, as well as 11% (1384 compared to 1250) glyco-peptides, 18% (1351 compared to 1141) site-specific glycans, and 18% (229 compared to 194) glycosites than FragPipe (Figure 1c and Figure S3c). Taken together, GlycanFinder 2.0 was chosen as the most suitable data analysis solution for glycoproteome characterization of the clinical cohort in this study.

**Figure 1.**
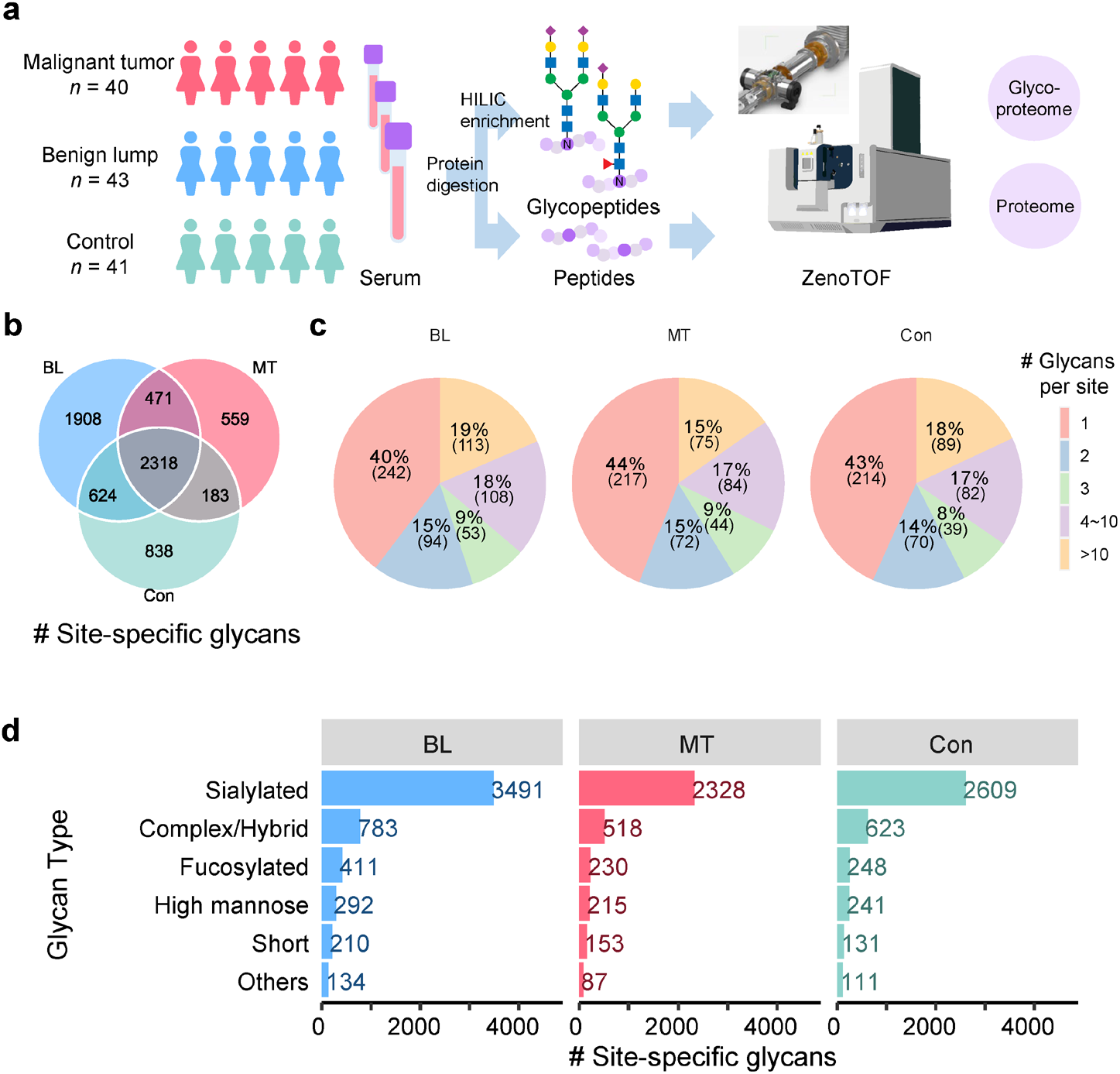
Glycoproteome profiling of the clinical serum samples. (**a**) Schematic illustration of the workflow where sera of breast malignant tumor (MT) patients, breast benign lump (BL) patients, and non-disease controls (Con) were subjected to proteomic and glycoproteomic analyses using ZenoTOF. (**b**) Overlap of the identified site-specific glycans from the glycoproteome samples of MT, BL and Con. (**c**) Distribution of protein glycosites attached with different number of unique glycans. (**d**) Distribution of glycan types of the identified site-specific glycans.

### Profiling serum glycoproteome for breast cancer

We analyzed serum samples from 124 female participants, including 40 breast malignant tumor (MT) patients, 43 breast benign lump (BL) patients, and 41 controls (Con) without breast diseases (**Table S1**). For each sample, proteomic and glycoproteomic analyses were performed (**Figure 1a**). At the proteome level, totally 1188 proteins were identified from the samples, among which 97% were shared by MT, BL, and Con (**Figure S2a**). The glycoproteome presented higher diversity than the proteome. From MT, BL, and Con, 3531, 5321, and 3963 site-specific glycans were identified, corresponding to 492, 610, and 494 glycosites, respectively, with 2318 site-specific glycans and 333 glycosites shared among them (**Figure 1b** and **Figure S2b**). Comparing glycoproteome identifications, the number of site-specific glycans identified from BL were higher than Con, whereas those from MT were lower than Con.

Intact glycopeptide analysis enables the characterization of system-wide glycosylation patterns. The identified glycosites showed expected N–X–S/T and N–X–C (less abundant) sequons (**Figure S2c**). We observed that 32%–37% of glycosites contained more than 3 linked glycans, 22%–24% contained 2 or 3 glycans, and 40%–44% had only 1 glycan (**Figure 1c**).

The overall proportions of different types of site-specific glycans in MT and BL were comparable to those of Con (**Figure 1d** and **Figure S3**). Sialylation was the most prominent (65%) glycosylation modifications. Non-sialylated and non-fucosylated complex or hybrid glycans accounted for 15%–16% of the site-specific glycans. Lower proportions of fucosylation (6%–7%) and high-mannose glycans (5%–6%) were observed.

### More significant alteration of glycoproteome than proteome in breast cancer

We then explored the quantitative profiles of the proteome and glycoproteome in MT, BL, and Con. Based on the glyco-proteome identification result, label-free quantification (LFQ) was conducted in the GlycanFinder workflow with match-between-runs, where glycopeptides identified by MS/MS in one run could be transferred to another according to their precursor ion *m*/*z* and LC retention time when applicable^33^. This mechanism can reduce missing values originated from stochasticity in MS/MS acquisition. Principal component analysis (PCA) was performed using the quantities of proteins and site-specific glycans (**Figure 2a**). The PCA score plot using the site-specific glycans showed an obvious separation among MT, BL, and Con within the first three components, while no obvious difference was observed using the protein quantities.

**Figure 2.**
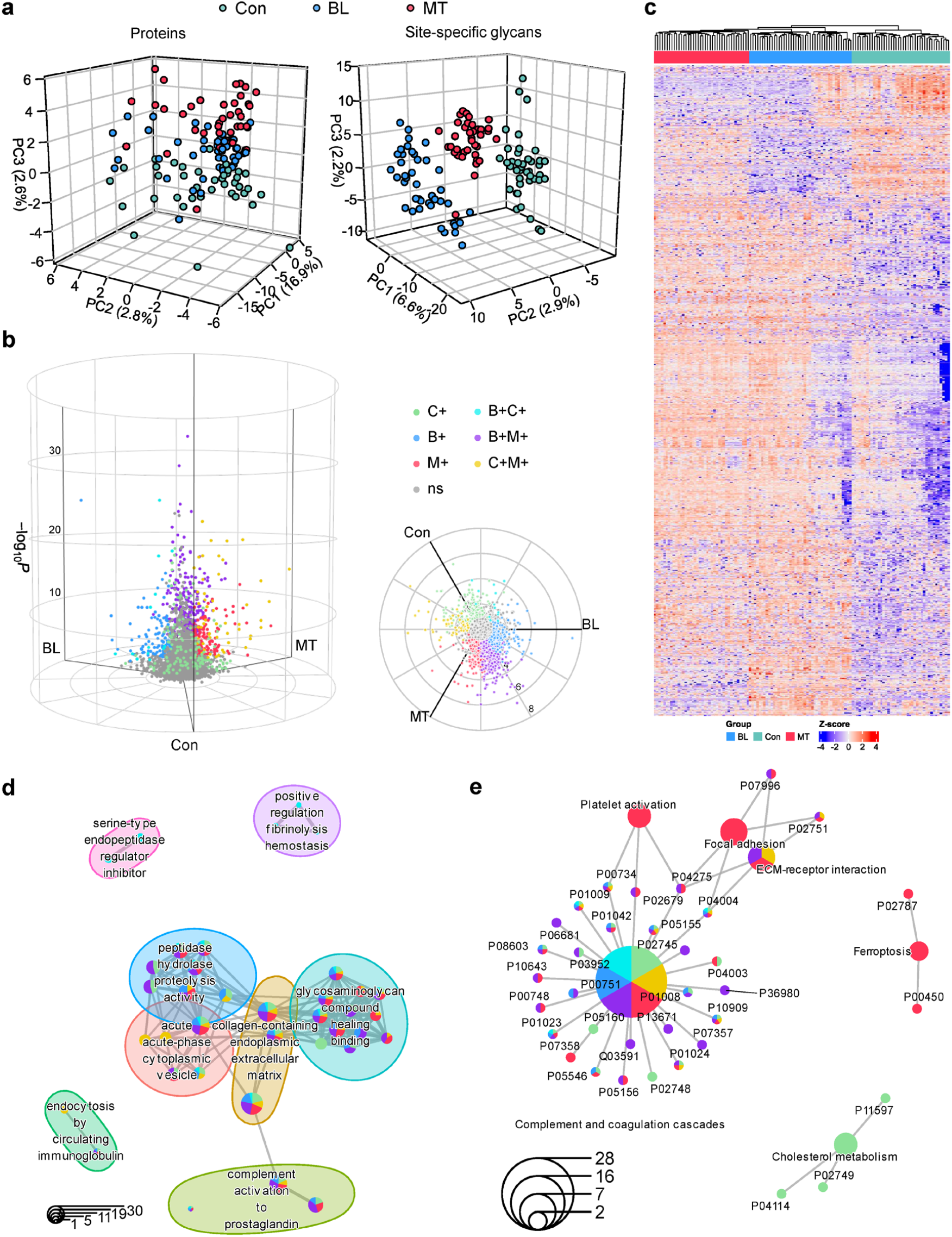
Quantitative alteration of serum glycoproteome in breast diseases. (**a**) Principal-component analysis (PCA) using the quantified site-specific glycans from the glycoproteome samples, compared with the quantified proteins from the proteome samples. (**b**) The 3D volcano plot showing differential site-specific glycans among MT, BL and Con. In the cylindrical coordinates, the radial axis shows fold change, the azimuthal angle conveys the degree to which a site-specific glycan is associated with one or more group, and the vertical axis shows −log_10_ adjusted p-value for three-way likelihood ratio test. An adjusted p-value < 0.01 for likelihood ratio test and a log_2_ fold change > 1 in radial scale are necessary conditions for a site-specific glycan considered significant. Colors demonstrate pairwise comparisons (adjusted p-value < 0.01 by moderate t-test) between the 3 groups: primary colors denote upregulation in one group only (red for MT, blue for BL, and green for Con) compared to reference group with minimum protein expression; composite colors show proteins significantly upregulated in two groups (purple for MT and BL, yellow for MT and Con, as well as cyan for BL and Con). Non-significant site-specific glycans (ns) are colored gray. A top view is shown. (**c**) Heatmap showing the hierarchical clustering of the samples using the differential site-specific glycans. (**d**) GO enrichment analysis of the differential site-specific glycans. The GO terms are clustered based on their similarities. Colors of the pie plots indicate the proportion of glycoproteins upregulated in different groups in each GO term. The rule of color is same as that in (**b**). (**e**) KEGG enrichment analysis of the differential site-specific glycans. KEGG pathways and proteins are present in a network. Colors of the pie plots indicate the occurrence of glycoproteins in each KEGG pathway or site-specific glycans from each glycoprotein upregulated in different groups. The rule of color is same as that in (**b**).

Differentially expressed site-specific glycans were determined by likelihood ratio test (adjusted p-value < 0.01), pair-wise moderated t-test (adjusted p-value < 0.01), and radical-scale fold change (absolute log_2_ fold change > 1). The three-way volcano plots^24^ (**Figure 2b**) demonstrated that the largest groups of differential site-specific glycans were upregulated in both MT and BL (345) or MT alone (230). A smaller number of site-specific glycans were uniquely upregulated in either BL (162) or Con (126), with only a few associated with them both (13).

The differential site-specific glycans were subjected to unsupervised hierarchical clustering (**Figure 2c**). The MT samples formed a cluster. The BL samples were separated into 2 parts, with 60% closer to the MT cluster and the remaining 40% farther away. All the Con samples assembled a unique cluster independent of MT and BL. In comparison, only a few differentially expressed proteins were found with the same statistical criteria (**Figure S4a**). The three groups of samples were poorly clustered using the differential proteins (**Figure S4b**). These results suggested that profiling the serum glycoproteome would be more informative for the diagnosis of breast cancer than measuring the serum proteome alone.

To investigate the functional characteristics of the differential site-specific glycans, gene ontology (GO)^34^ enrichment analysis was performed using their corresponding glycoproteins. The enriched GO terms were grouped into 8 clusters based on their similarities (**Figure 2d** and **Figure S5**). The largest cluster was related to coagulation and glycosaminoglycan binding. Other clusters involved peptidase activity, immune response, vesicle lumen, and extracellular matrix. Notably, site-specific glycans on the same glycoprotein may vary in expression patterns and be upregulated in different groups. The glycoproteins of the differential site-specific glycans were further mapped to metabolic pathways in the Kyoto Encyclopedia of Genes and Genomes (KEGG)^35^ database (**Figure 2e** and **Figure S6**). Among the enriched KEGG pathways, complement and coagulation cascades were substantially altered in MT and BL with quite a few site-specific glycans upregulated in different groups. Cholesterol metabolism was downregulated in MT and BL, while focal adhesion and ferroptosis were upregulated in MT.

### Discovering glyco-signatures for breast cancer diagnosis

To screen potential biomarkers from the site-specific glycans that contributed to breast cancer diagnosis, we used random forest, a widely-used machine learning algorithm, to build a sample classifier (**Figure 3a**). Advantages of random forest include that it allows the efficient classification using large number of features (site-specific glycans in this study) without dimension reduction in advance and the estimation of the importance of each feature. The cohort were randomly split into the modeling subset (25 MT samples, 28 BL samples, and 26 Con samples) and the holdout subset (15 samples for each group). In the context of cancer diagnosis, BL and Con samples were labeled as non-MT, so that the task was a binary classification problem. The modeling subset was first used for feature selection by Monte-Carlo cross validation (MCCV). In each MCCV, 2/3 of the samples were used to evaluate the feature importance and the classification model built using the important features was validated on the remaining 1/3 samples. Receiver operating characteristic (ROC) curves were used to evaluate the discrimination power of the classification models built using the important features (**Figure 3b**). The number of important features was surveyed. Using the 5-to 50-feature models, the area under the curve (AUC) values were 0.89 to 0.96, and the 15-feature models achieved a trade-off between accuracy (AUC 0.94) and model complexity. The features were ranked by their frequencies of being selected in 15feature models during MCCV (**Figure 3c**).

**Figure 3.**
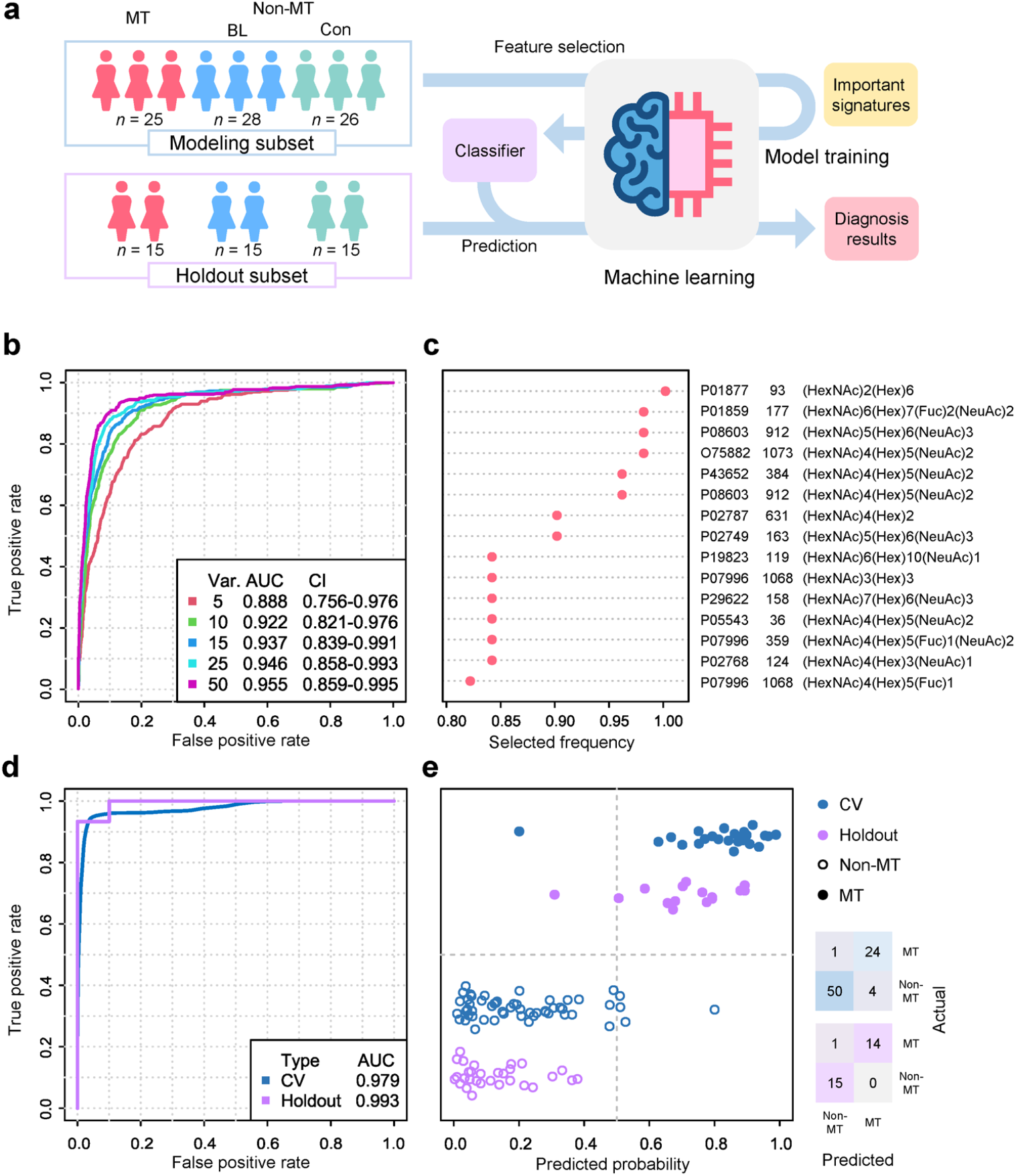
Site-specific glycan biomarker screening for breast cancer diagnosis. (**a**) Schematic illustration of the machine learning workflow for biomarker screening. (**b**) ROC curves of models using different numbers of important features by cross validation (CV) on the modeling subset. AUC values and confidence intervals (CI) are indicated. (**c**) Features ranked by the frequency of being selected in the 15-feature models during CV. The monosaccharide symbols are defined in **Table S2**. (**d**) ROC curves of the classifier using the 15 mostly selected features, evaluated by CV on the modeling subset and tested on the holdout subset. (**e**) Prediction accuracy of the classifier. Confusion matrices are present for the modeling and holdout subsets.

The 15 mostly selected features were used to train a classifier with the modeling subset by MCCV, resulting in an AUC of 0.98 (**Figure 3d**). With a predicted probability threshold of 0.5, 96% (24/25, recall) of the MT samples were correctly detected, accounting for 86% (24/28, precision) of the predicted positive cases (**Figure 3e**). The classifier was then tested on the hold-out subset, achieving an AUC of 0.99, recall of 93% (14/15), and precision of 100% (15/15).

We further determined the AUC for the individual signatures (**Figure 4, Figure S7** and **S8**). Most of these signatures yielded favorable performance with AUC values greater than 0.87 in the modeling subset and 0.81 in the holdout subset. Hence, these site-specific glycans could be considered as biomarker candidates for breast cancer diagnosis.

**Figure 4.**
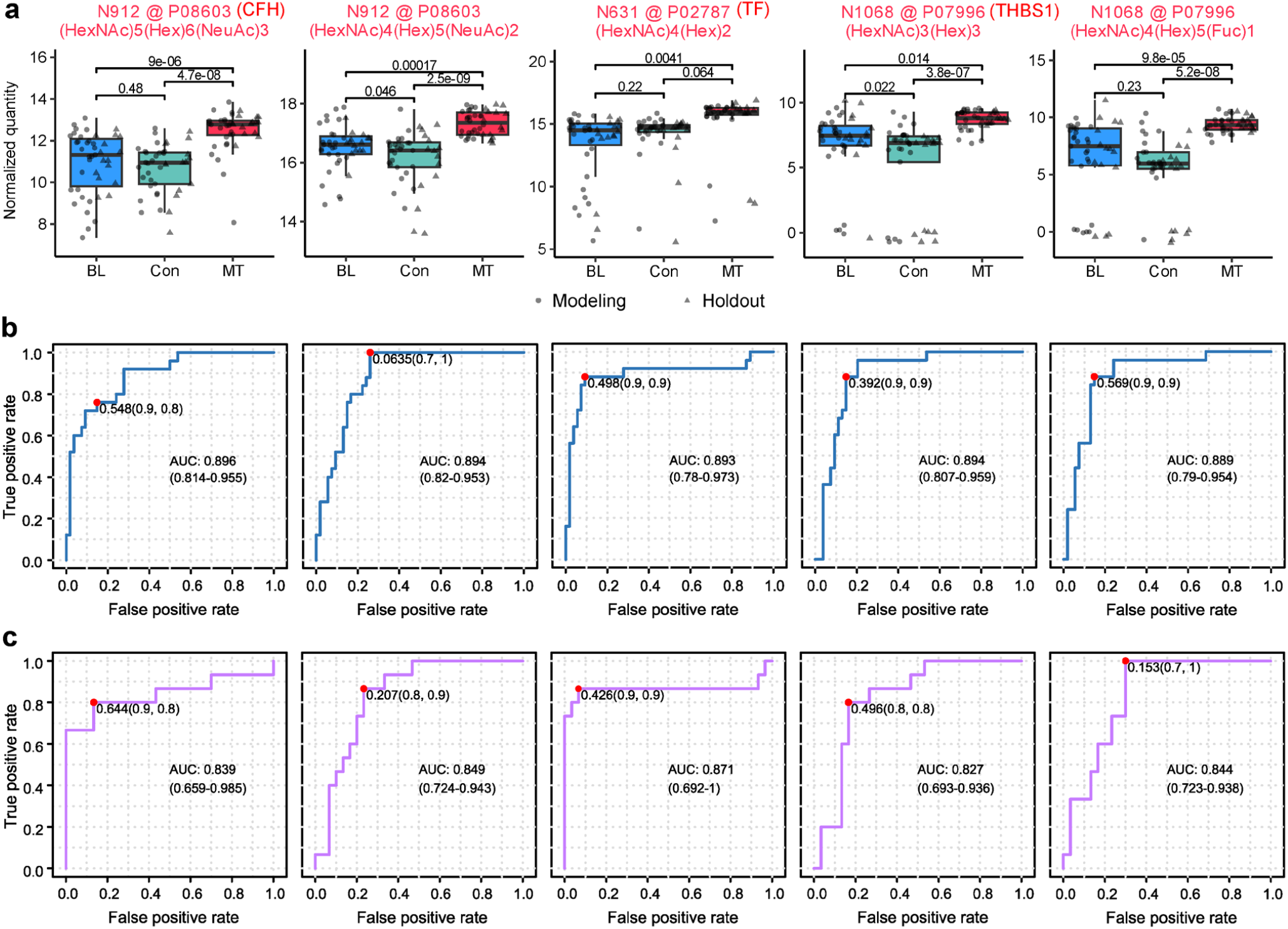
Site-specific glycans considered as glyco-signatures for breast cancer diagnosis. (**a**) Quantities of the site-specific glycans in MT, BL, and Con. Adjusted p-values of pairwise moderated t-test are indicated. Red text color demonstrates up-regulations in MT (adjusted p-value < 0.01 and log_2_ fold change > 1). (**b**) ROC curves for the samples in the modeling subset. (**c**) ROC curves for the samples in the holdout subset. AUC values and confidence intervals (CI) are indicated. Red dots represent the optimal cutoff values. Other glyco-signatures are present in **Figure S7** and **S8**. The monosaccharide symbols are defined in **Table S2**.

## DISCUSSION

In this work, we developed a method for intact glycopeptide characterization with ZenoTOF and demonstrated its ability on serum glycoproteomics for potential biomarker discovery. While most of the glycoproteomics studies used Orbitrap-based platforms, we optimized the CID parameters, achieving a precursor *m*/*z*–dependent, dynamic and stepped CE setting that were similar to the Orbitrap-specific HCD for intact glycopeptide fragmentation. We also found the most suitable glycoproteomic informatics solution for ZenoTOF and assessed the performance of other community-available tools. Typical Q-TOF systems can provide higher scan rates and do not suffer from space-charge effects restricting usable ions like trap-based systems^36^. The Zeno trap can align the time of ions arriving in the TOF accelerator with the pulse to the TOF, resulting in higher duty cycle and greatly improves sensitivity as reported in recent proteomics studies^18, 37^. In addition, our method adopted the microflow LC system instead of nanoflow LC routinely used in proteomic studies. Previous research has demonstrated that microflow LC provided improved system stability benefiting the precise quantification of glycopeptides^14^. This feature would facilitate extensive data acquisitions for large-scale projects and in diagnostic laboratories. We present the complete workflow of MS acquisition and data analysis for proteome-wide and site-specific profiling of glycosylation using ZenoTOF, which provides an alternative solution to expand the toolbox of glycoproteomics.

Breast cancer is the most common cancer and one of the leading causes of cancer death in women^38^. Serum or plasma biomarkers are especially appealing for the potential to detect cancer in early stages that are increasingly treatable. Although candidates have been suggested as diagnostic markers of breast cancer, including carcinoembryonic antigen, cancer antigen 15-3, and cancer antigen 27-29, they lack sensitivity and/or specificity for early disease detection^39^. For biomarker discovery, previous studies have profiled glycoproteomes^40, 41^ and glycomes^42^ where the glycans were chemically cleaved from glycosylated proteins. We employed our method to analyze the intact glycopeptides from clinical serum samples and quantified the site-specific glycosylation changes in breast diseases. Much more significant changes of site-specific glycans were observed across the sample groups than those at the protein level, indicating that serum glycoproteome could be a better resource for biomarker discovery compared to the proteome. Using machine learning, we selected 15 site-specific glycans to construct a classifier model for breast cancer detection, achieving a high accuracy validated on the holdout samples. Most of the individual site-specific glycans yielded AUC values over 0.8, demonstrating their powerful prediction capability as glyco-signatures for the detection of breast cancer.

The 15 site-specific glycans used in the classifier are from 12 glycoproteins (**Figure 4, Figure S7** and **S8**). Notably, most of these glycoproteins were reported to be relevant to breast cancer. Complement factor H (CFH, UniProt accession P08603) is a central regulator of the complement system. It has been reported that CFH expressed by human breast cancer cells correlates with immunosuppression, breast cancer recurrence and severity of the disease^43^. We observed a significant up-regulation of the glycans (HexNAc)5(Hex)6(NeuAc)3 and (HexNAc)4(Hex)5(NeuAc)2 at the glycosite N912 in CFH. Serotransferrin (TF, UniProt accession P02787) is an iron binding transport protein related to ferroptosis. A previous study on serum transcriptome and proteome has reported the down-regulation of TF in breast cancer^44^. We did not observe an alteration of TF at the protein level between MT and Con, but an up-regulation of the glycan (HexNAc)4(Hex)2 at the glycosite N631. Thrombospondin-1 (THBS1, UniProt accession P07996) is a component of the extracellular matrix with roles in regulating cancer development. High level of THBS1 has been reported in breast cancer plasma^45^. We observed that the glycans (HexNAc)3(Hex)3 and (HexNAc)4(Hex)5(Fuc)1 at the glycosite N1068 in THBS1 were up-regulated in MT. Other selected features included glycans from the immunoglobulin heavy constant regions alpha 2 (IGHA2, UniProt accession P01877) and gamma 2 (IGHG2, UniProt accession P01859), attractin (ATRN, UniProt accession O75882), afamin (AFM, UniProt accession P43652), and beta-2-glycoprotein 1 (APOH, UniProt accession P02749). IGHA2 expression was consistently positive in tumor cells that were metastatic to regional nodes^46^. AFM was found to be higher in sera of prediagnostic breast cancer^47^. APOH was increased in sera of the breast cancer patients^48^. We found site-specific glycans from these glycoproteins up-regulated in MT compared to Con, while regulations at the protein level (**Figure S9**) were not significant or with smaller fold change (log_2_FC < 1). In regard to the divergence of regulations between the site-specific glycans and proteins, the detailed roles of these site-specific glycans in the pathogenesis of breast cancer require further investigation.

## CONCLUSIONS

In summary, we present a method for intact glycopeptide characterization using ZenoTOF and showcase its use for large-scale site-specific quantitative glycoproteome profiling of clinical serum samples. Our findings highlight that sitespecific glycoproteome provides exciting opportunities for disease biomarker study. We expect that the method will have promising potential in basic and clinical glycoproteomic studies.

## AUTHOR INFORMATION

### Author Contributions

Yi Yang: Conceptualization; Funding acquisition; Investigation; Writing – original draft. Dan Zhao: Investigation. Ji Luo: Investigation. Yuxiang Lin: Resources; Ling Lin: Investigation; Baozhen Shan: Investigation; Software. Hongxu Chen: Investigation; Supervision. Liang Qiao: Conceptualization; Funding acquisition; Supervision; Writing – review & editing.

\# These authors contributed equally: Yi Yang, Dan Zhao, and Ji Luo.

### Notes

J.L. and H.C. are employed by SCIEX. B.S. is employed by Bioinformatics Solutions Inc. The other authors declare no conflict of interest.

## ACKNOWLEDGMENT

This work was supported by Zhejiang Provincial Natural Science Foundation of China (LQ24B050003), the Science and Technology Commission of Shanghai Municipality (23JS1400100), and the National Natural Science Foundation of China (NSFC, 22374031).

## Table of Contents (TOC) artwork

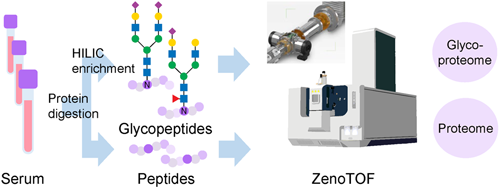

