## Supporting Information for "Quantitative site-specific glycoproteomics by ZenoTOF reveals glyco-signatures for breast cancer diagnosis"

Yang et al.

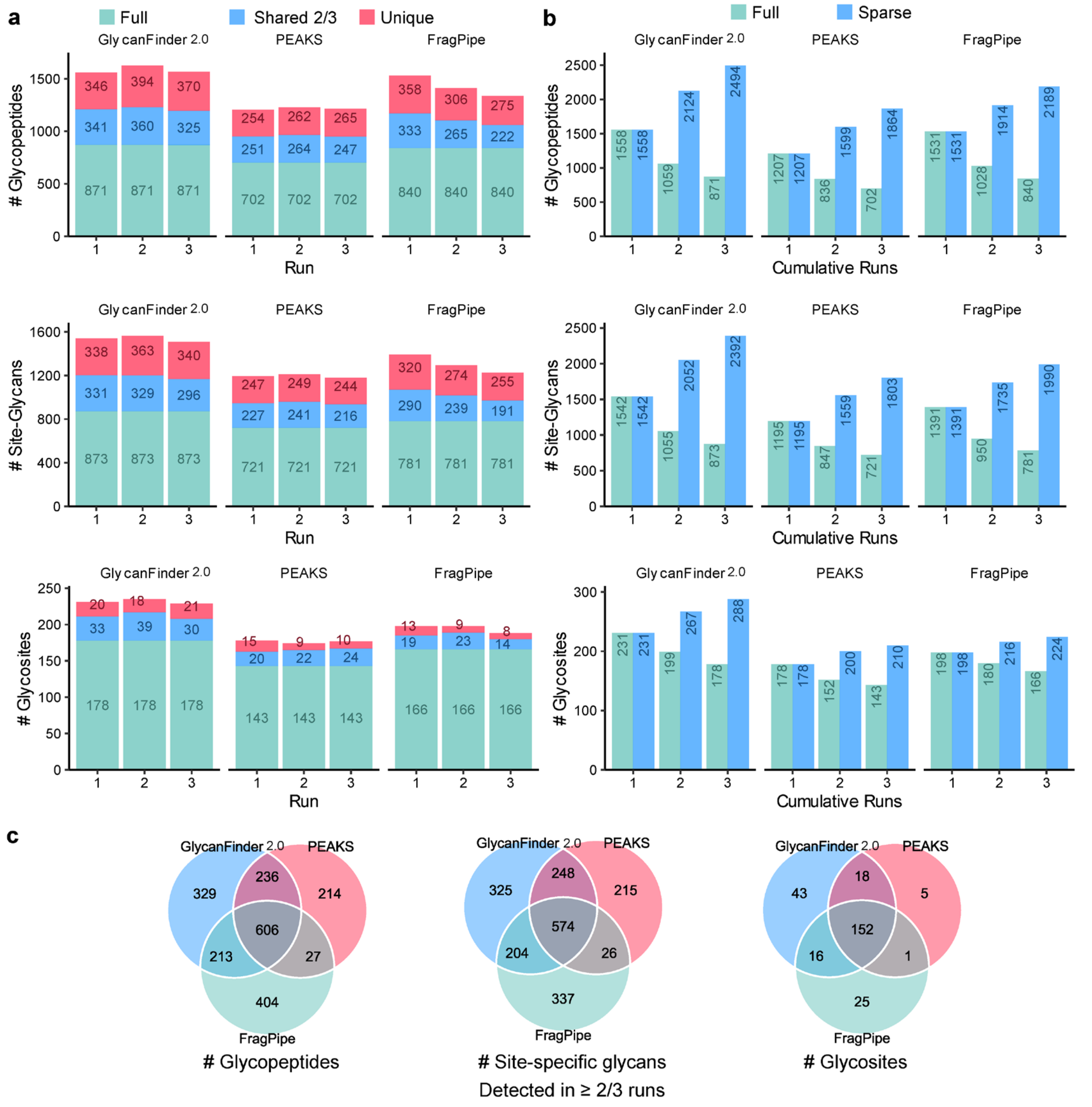


**Figure S1.** Identification results of the pooled serum sample at the glycopeptide, site-specific glycan, and protein glycosite level using different software.

(**a**) Numbers of identifications per run. "Full" represents identifications observed in all the runs; "shared 2/3" represents identifications observed in 2 runs; "unique" represents identifications observed in only 1 run. (**b**) Numbers of cumulative identifications from run 1 to 3. "Full" represents identifications shared in all the cumulative runs; "sparse" represents identifications observed in at least one run in the cumulative runs. (**c**) Comparison of numbers of identifications shared in ≥2/3 runs.

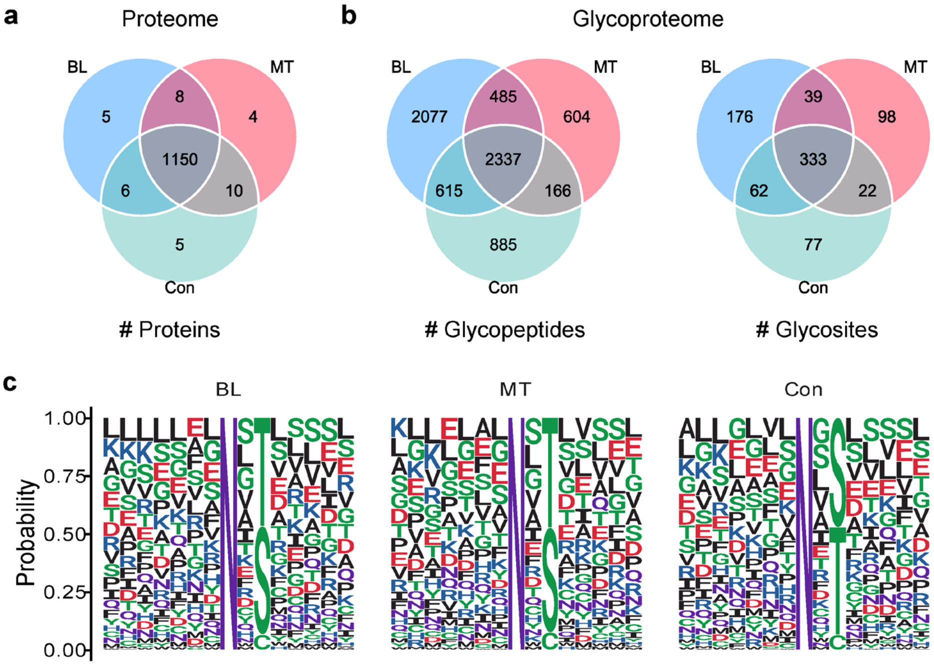


**Figure S2.** Identification results of the clinical serum samples.

(**a**) Overlap of the identified proteins from the proteome samples of MT, BL and Con. (**b**) Overlap of the identified glycopeptides and protein glycosites from the glycoproteome samples of MT, BL and Con. Related to **Fig. 1b**. (**c**) Sequence motifs distribution for the identified glycosites.

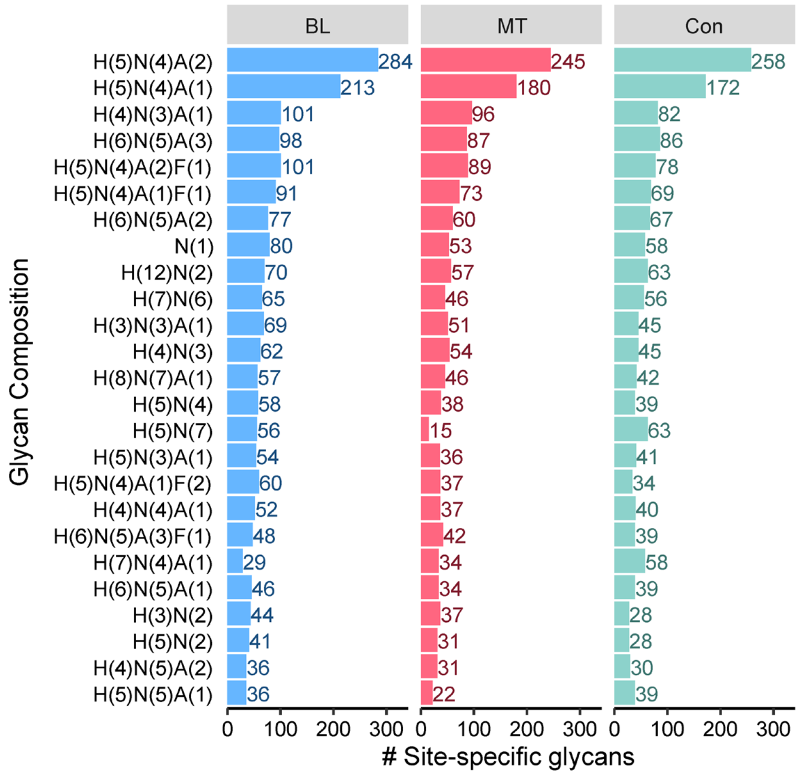


**Figure S3.** Distribution of glycan composition of the identified site-specific glycans. Related to **Fig. 1d**. The monosaccharide symbols are defined in **Table S2**.

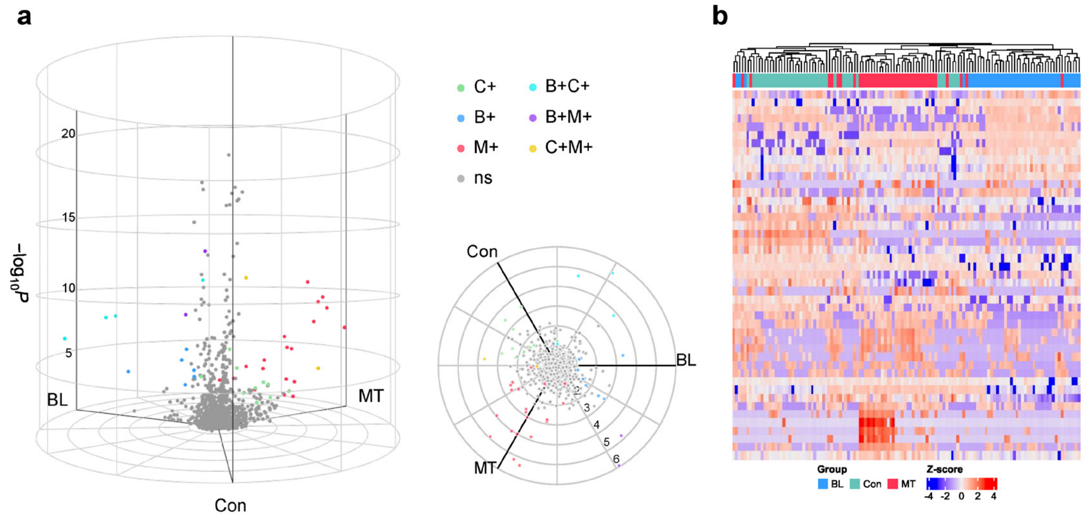


**Figure S4.** Differentially expressed proteins from the clinical serum proteome samples.

(**a**) The 3D volcano plot showing differential proteins among MT, BL and Con. In the cylindrical coordinates, the radial axis shows fold change, the azimuthal angle conveys the degree to which a protein is associated with one or more group, and the vertical axis shows $-\log_{10}$ adjusted p-value for three-way likelihood ratio test. An adjusted p-value $< 0.01$ for likelihood ratio test and a $\log_2$ fold change $> 1$ in radial scale are necessary conditions for a protein considered significant. Colors demonstrate pairwise comparisons (adjusted p-value $< 0.01$ by moderate t-test) between the 3 groups: primary colors denote upregulation in one group only (red for MT, blue for BL, and green for Con) compared to reference group with minimum protein expression; composite colors show proteins significantly upregulated in two groups (purple for MT and BL, yellow for MT and Con, as well as cyan for BL and Con). Non-significant proteins (ns) are colored gray. A top view is shown. (**b**) Heatmap showing the hierarchical clustering of the samples using the differential proteins.

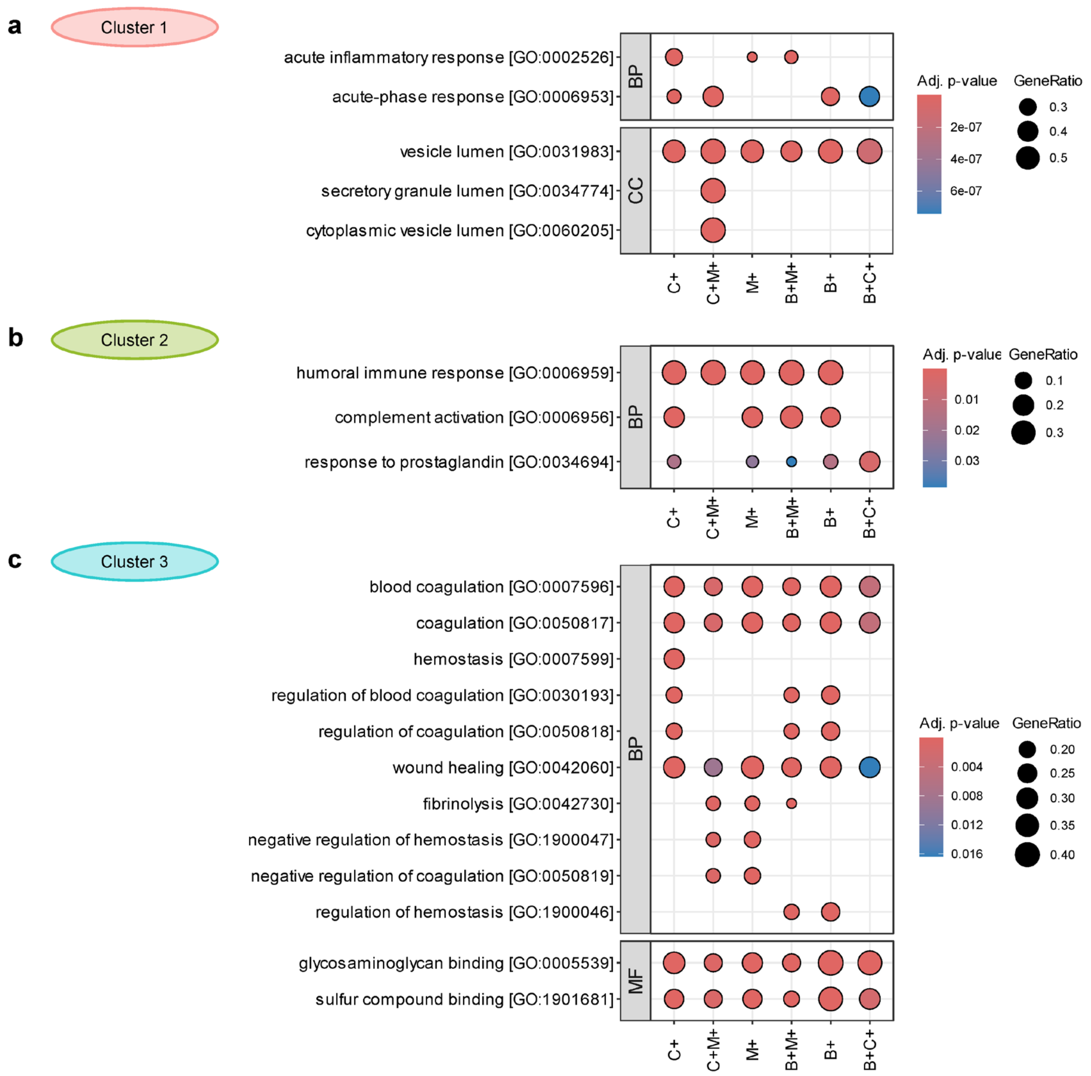


**Figure S5.** GO enrichment analysis of the differential site-specific glycans.

(**a**) Enriched GO terms related to the keywords “acute”, “acute-phase”, “cytoplasmic”, and “vesicle” (cluster 1). (**b**) Terms related to “complement”, “activation”, and “prostaglandin” (cluster 2). (**c**) Terms related to “glycosaminoglycan”, “compound”, “healing”, and “binding” (cluster 3). (Continued on next page)

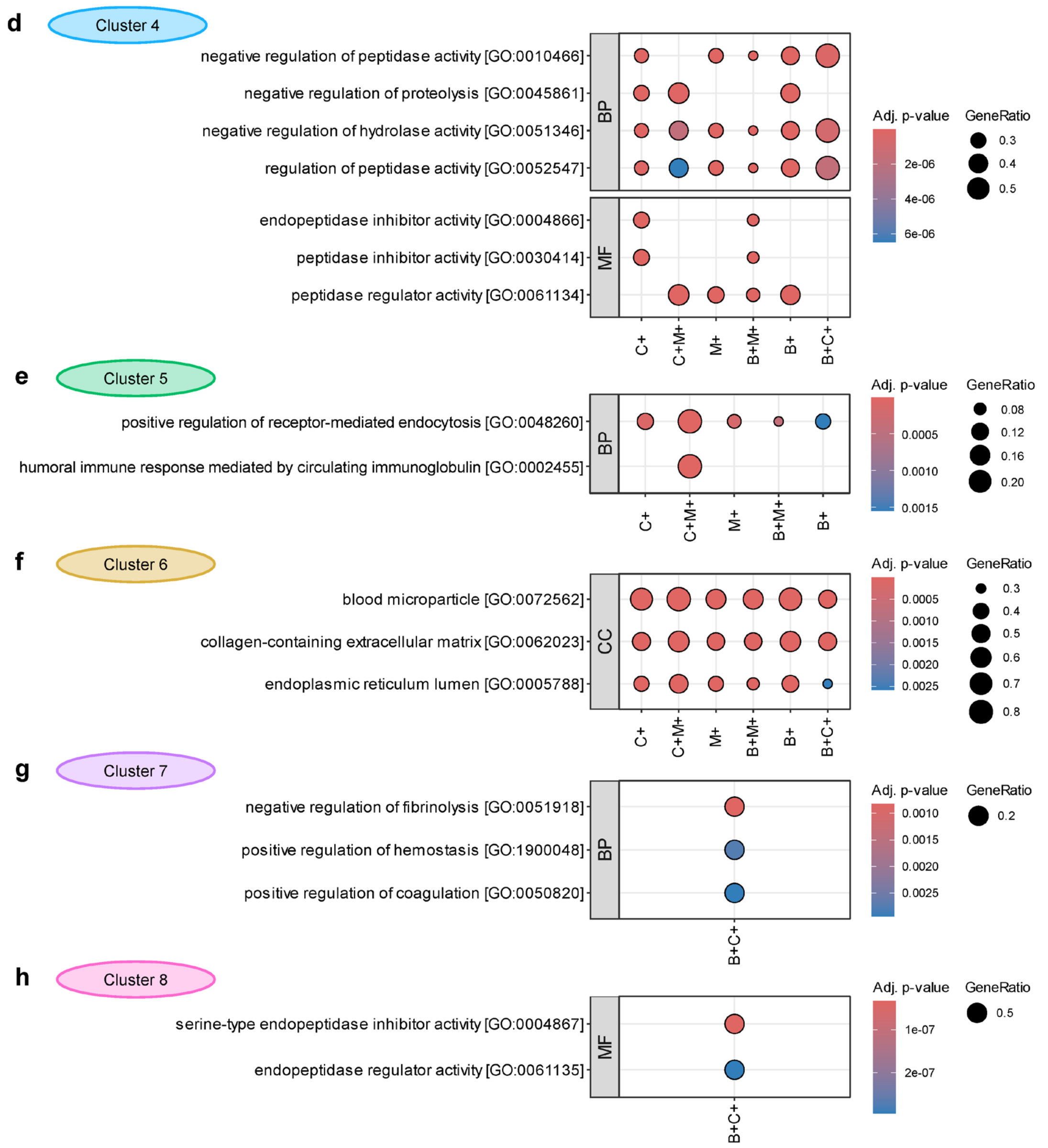


**Figure S5.** (Continued)
(**d**) Terms related to the keywords "peptidase", "hydrolase", "proteolysis", and "activate" (cluster 4). (**e**) Terms related to "endocytosis", "circulating", and "immunoglobulin" (cluster 5). (**f**) Terms related to "collagen-containing", "endoplasmic", "extracellular", and "matrix" (cluster 6). (**g**) Terms related to "positive", "regulation", "fibrinolysis", and "hemostasis" (cluster 7). (**h**) Terms related to "serine-type", "endopeptidase", "regulator", and "inhibitor" (cluster 8). Related to **Fig. 2d**, which shows the clustering of the terms. Upregulation in one group only (M+ for MT, B+ for BL, and C+ for Con) and composite upregulation in two groups (B+M+, C+M+, and B+C+) are abbreviated to initials.

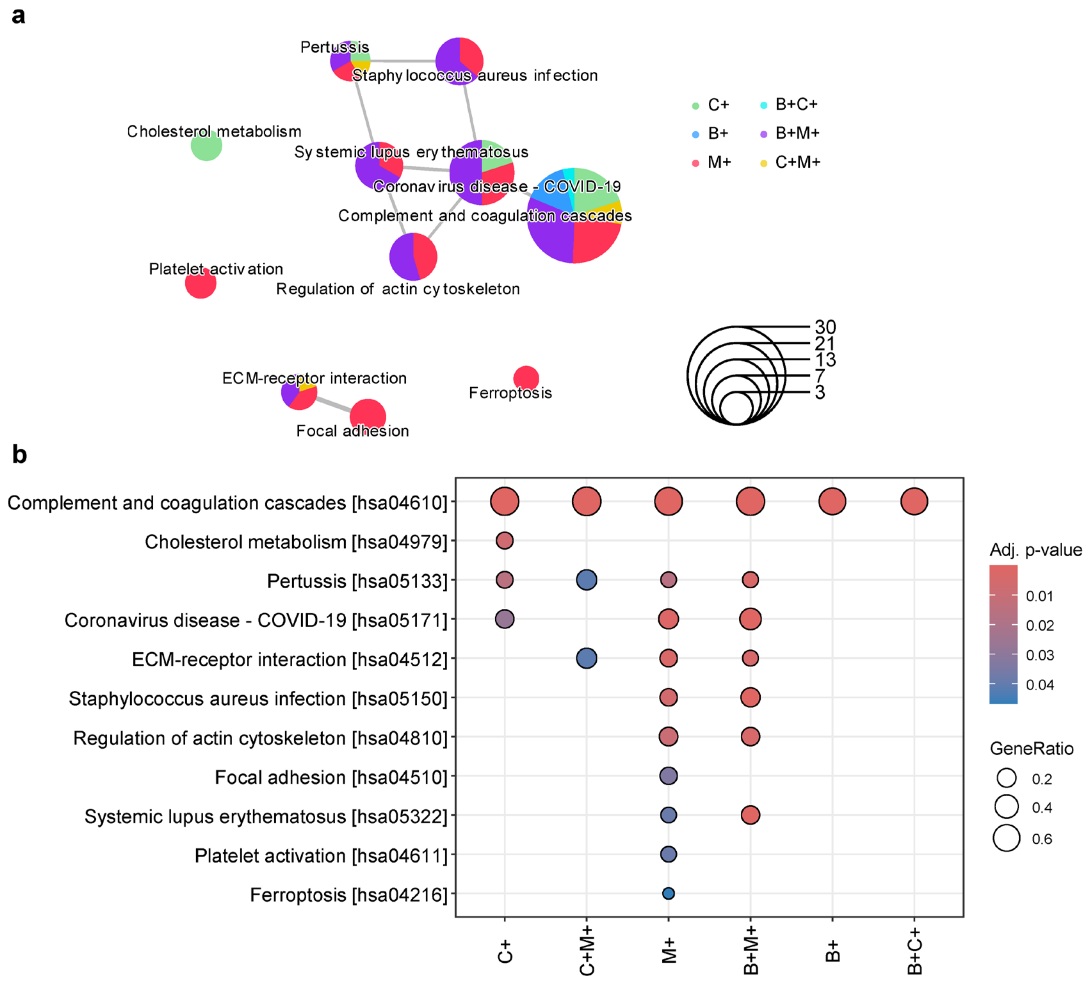


**Figure S5.** KEGG enrichment analysis of the differential site-specific glycans.

(**a**) Network of the enriched KEGG pathways. Colors of the pie plots indicate the proportion of glycoproteins upregulated in different groups in the pathway. The rule of color is same as that in **Fig. 2b**. (**b**) Enriched KEGG terms. Related to **Fig. 2e**. Upregulation in one group only (M+ for MT, B+ for BL, and C+ for Con) and composite upregulation in two groups (B+M+, C+M+, and B+C+) are abbreviated to initials.

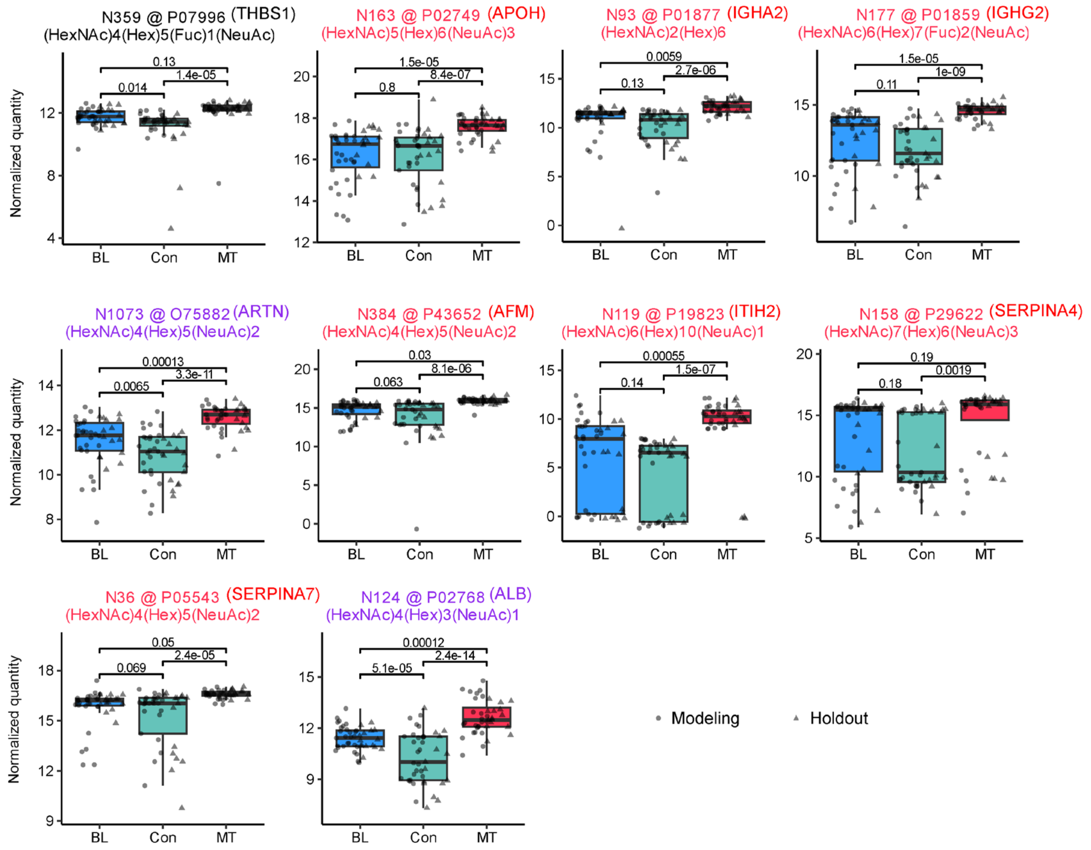


**Figure S7.** Quantities of site-specific glycans considered as glyco-signatures for breast cancer diagnosis.

Adjusted p-values of pairwise moderated t-test are indicated. Text colors demonstrate up-regulations (adjusted p-value < 0.01 and $\log_2$ fold change > 1; red for MT, purple for MT and BL, as well as black for no significant change. The monosaccharide symbols are defined in **Table S2**.

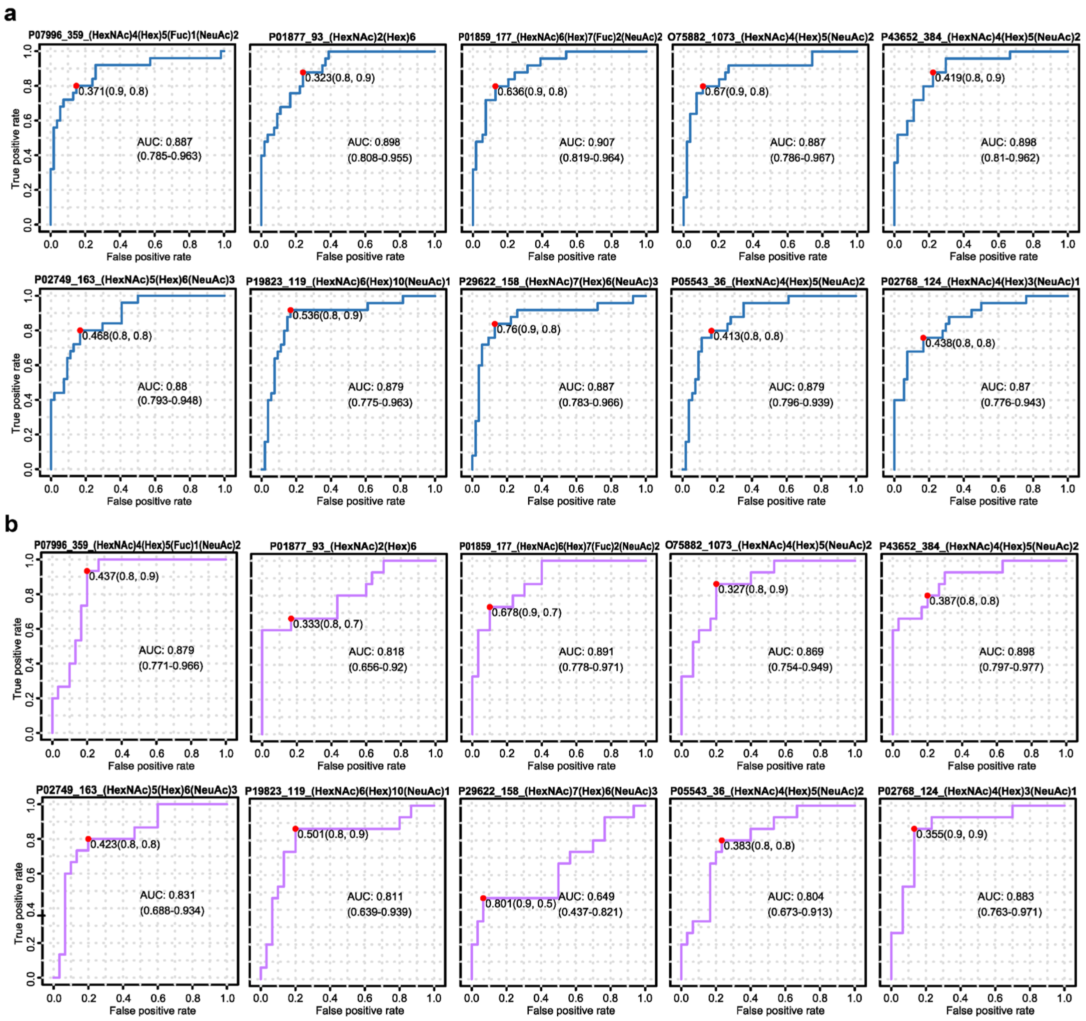


**Figure S8.** ROC curves of site-specific glycans considered as glyco-signatures for breast cancer diagnosis.

(**a**) ROC curves for the samples in the modeling subset. (**b**) ROC curves for the samples in the holdout subset. AUC values and confidence intervals (CI) are indicated. Red dots represent the optimal cutoff values. The monosaccharide symbols are defined in **Table S2**.

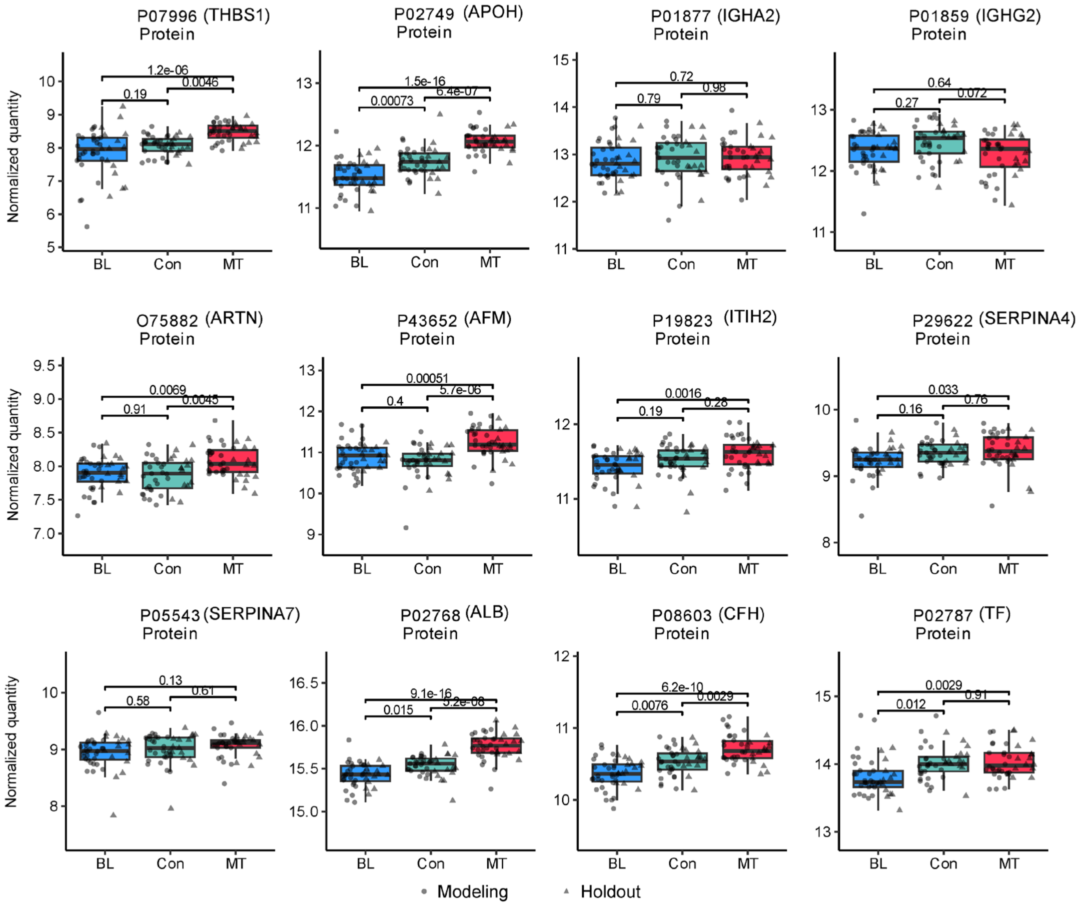


**Figure S9.** Quantities of proteins detected from the proteome samples corresponding to the 15 site-specific glycans considered as glyco-signatures for breast cancer diagnosis.

Related to **Figure 4** and **Figure S7**. Adjusted p-values of pairwise moderated t-test are indicated. Text colors demonstrate up-regulations (adjusted p-value < 0.01 and $\log_2$ fold change > 1; red for MT, purple for MT and BL, as well as black for no significant change. The monosaccharide symbols are defined in **Table S2**.

**Table S1. Demographic data of the breast disease patients and normal controls.**

| Characteristics | Malignant tumor patients (MT) | Benign lump patients (BL) | Normal controls (Con) |
|---|---|---|---|
| Number | 40 | 43 | 41 |
| Sex | All are women | | |
| Age (year, mean ± SD) | 50.48 ± 9.98 | 43.95 ± 8.89 | 48.73 ± 9.78 |

SD: standard deviation.

**Table S2.** Nomenclature of monosaccharides involved in this study.

| Code | Abbr. | Symbol | Description |
|---|---|---|---|
| H | Hex | 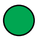<br>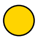 | Hexose<br>Mannose<br>Galactose |
| N | HexNAc | 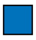<br>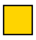 | *N*-Acetylhexosamine<br>*N*-Acetylglucosamine (GlcNAc)<br>*N*-Acetylgalactosamine (GalNAc) |
| A | NeuAc | 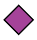 | *N*-Acetylneuraminic acid |
| F | Fuc | 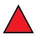 | Fucose |

**Method S1.** Sample preparation and MS analysis.

**Glycopeptide sample preparation**

Individual human sera of 10 μL were added to 90 μL of 8 M urine to denature the proteins. The proteins were then reduced with 70 mM tris(2-carboxyethyl)phosphine at 37 °C for 1 h and alkylated with 140 mM 2-chloroacetamide for 0.5 h at room temperature in the dark. After diluted with 100 mM triethylammonium bicarbonate buffer, trypsin was added to reach a final enzyme-to-substrate ratio of 1:50 (wt/wt) and incubated at 37 °C overnight. The reaction was terminated by adding 0.5% trifluoroacetic acid (TFA). All digested samples were centrifuged at 16,000 × g for 10 min and the supernatants were desalted using the Sep-Pak C18 cartridges (Waters, USA). The desalted peptides were then dried by vacuum centrifugation and used for glycopeptide enrichment.

Glycopeptides were enriched using hydrophilic interaction liquid chromatography (HILIC). The desalted peptides (500 μg) were resuspended in 300 μL loading buffer containing 80% acetonitrile and 1% TFA and then loaded onto an in-house micro-column containing 50 mg of ZIC-HILIC particles (Merck Millipore, Germany) packed onto a C8 disk. The flow through was collected and reloaded onto the column for additional four times. Then, the column was washed with 200 μL loading buffer for four times. Finally, the glycopeptides that have been enriched in the column were collected by elution with 600 μL 0.1% TFA and dried by vacuum centrifugation.

**LC-MS/MS analysis**

The enriched glycopeptides from the clinical samples were resuspended in 0.1% FA and submitted for microflow LC-MS/MS analysis, which was performed in a Microscale LC System (Waters, USA) coupled online to a ZenoTOF 7600 mass spectrometer (SCIEX, USA). Glycopeptides (8 μL injection volume, corresponding to 2 μg per sample) were loaded and separated on a Kinetex XB-C18 LC column (2.6 μm particle size, 100 Å pore size, 0.3 mm inner diameter × 150 mm, catalog number 00F-4496-AC, Phenomenex, China) at a flow rate of 5 μL $min^{-1}$. The column temperature was maintained at 45°C. Solvent A was a 0.1% FA aqueous solution. Solvent B was 80% acetonitrile containing 0.1% FA. The overall 60-min LC gradient was described as follows: initial at 2% B, from 2% to 5% B for 2 min, from 5% to 32% B for 41 min, from 32% to 55% B for 5 min, from 55% to 85% B for 0.5 min, held on 85% B for 4 min to clean the system, and finally back to 2% B in 0.5 min and for 7 min to equilibrate the system.

Data collection was performed by SCIEX OS (version 3.1.6.44). For glycoproteomic analysis, data-dependent acquisition (DDA, known as IDA on ZenoTOF) mode was operated to switch between MS and MS/MS acquisition. Ion source gas 1 and 2 were set to 16 and 35 psi, respectively; curtain gas was set to 35, CAD gas to 7, and source temperature to 250 °C; spray voltage was set to 5500 V. MS/MS scans were performed for the top 16 parent ions. Glycopeptide fragmentation was performed by CID (i.e., CAD on ZenoTOF) with dynamic CE. CES was set to 10.

For proteomic analysis, the peptides without enrichment (4 μL injection volume, corresponding to 1 μg per sample) were analyzed using the same LC-MS/MS instrument and the data-independent acquisition (DIA, known as SWATH MS on ZenoTOF) mode. A Zeno SWATH acquisition scheme with 85 isolation windows (variable window with a *m*/*z* overlap of 1) was used, covering a precursor *m*/*z* range of 399.5–904.5. Dynamic CE and CES of 10 were used. The other MS parameters were the same as those for glycopeptides.

For benchmarking purpose, a pooled glycopeptide sample (8 μL injection volume, corresponding to 2 μg per sample) was analyzed using different CID parameters and an overall 120-min LC gradient described as follows: initial at 2% B, from 2% to 4% B for 2 min, from 4% to 30% B for 91 min, from 30% to 52% B for 15 min, from 52% to 85% B for 0.5 min, held on 85% B for 5 min to clean the system, and finally back to 2% B in 0.5 min and for 6 min to equilibrate the system. Glycopeptide fragmentation was performed by CID with CE (center point) of 20, 30, and dynamic CE. CES was set to 10.

**Method S2.** Data analysis and bioinformatics.

**Glycoproteomics MS data analysis**

For the clinical glycoproteome samples, raw data (.wiff) were searched with GlycanFinder[1] (version 2.0, Bioinformatics Solutions Inc., Canada) against the UniProt *Homo sapiens* database (taxonomy ID 9606, reviewed, 20601 protein entries, access date 2022-09). Two missed cleavages were allowed for trypsin specific digestion. The fixed modification was carbamidomethylation of all cysteine residues (+57.02). Variable modifications included oxidation of methionine (+15.99) and acetylation on protein N-term (+42.01). For the benchmarking sample, PEAKS, and FragPipe[2, 3] were also used for glycopeptide identification.

**Proteomics MS data analysis**

Raw DIA data of the proteome samples were analyzed by Spectronaut[4] (version 18.4.231017, Biognosys AG, Switzerland) using the directDIA+ workflow. Q-value cut-off at both precursor and protein level (in the experiment context) was set as 1%. Automatic cross-run normalization was enabled. Other parameters were default.

**Bioinformatic analysis**

Bioinformatic analysis was conducted using R (version 4.3.1, https://www.r-project.org/). Differentially expressed site-specific glycans and proteins were determined using the R packages limma[5] (version 3.56.2) and volcano3D[6] (version 2.0.9) with the criteria of adjusted p-value < 0.01 by three-way likelihood ratio test and pairwise moderate t-test with Benjamini-Hochberg (BH) correction, and absolute log2 fold change > 1. GO and KEGG enrichment analysis was perform using the R package clusterProfiler[7] (version 4.8.3). Random forest machine learning was performed using the R package MetaboAnalystR[8] (version 4.0.0).

For data visualization, the following R packages were used: ggplot2 (version 3.4.3), VennDiagram[9] (version 1.7.3), ggseqlogo[10] (version 0.1), plot3D (version 1.4), ComplexHeatmap[11] (2.16.0), volcano3D[6] (version 2.0.9), and enrichplot (version 1.20.3).
